# Detection of nervous necrosis virus RGNNV genotype in pearl gentian grouper (*Epinephelus lanceolatus* ♂ × *E. fuscoguttatus* ♀) fry imported to Thailand

**DOI:** 10.1101/2022.02.24.481889

**Authors:** Sumit Vinod Jungi, Pakkakul Sangsuriya, Suwimon Taengphu, Kornsunee Phiwsaiya, Molruedee Sonthi, Bunlung Nuangsaeng, Krishna R Salin, Saengchan Senapin, Ha Thanh Dong

**Author notes:** Corresponding authors: S. Senapin, H.T. Dong.

## Abstract

Nervous necrosis virus (NNV) is a deadly virus that affects more than 120 fish species worldwide, but there is little information about it in Thailand. In August 2019, a population of pearl gentian grouper fry imported to Thailand experienced mass mortality. The diseased fish exhibited darkening, floating on the water surface, ‘sleepy behaviour’, and erratic swimming. We received a set of samples for disease diagnosis. PCR analysis revealed that these samples were positive for NNV but negative for Megalocytivirus ISKNV and Ranavirus. Sequencing the virus’s genome and phylogenetic analysis revealed that it is a member of the red-spotted grouper nervous necrosis virus (RGNNV) genotype. *In situ* hybridization using an NNV-specific probe revealed localization of the virus in the vacuolation lesions of the central nervous system (brain and spinal cord) and the retina of infected fish. The virus was successfully isolated from tissue homogenate from diseased fish using the E-11 cell line. This study identified NNV (RGNNV genotype) as a virus associated with massive die-offs in imported pearl grouper fry in Thailand. Considering the infectious nature of the virus and its broad host range, appropriate biosecurity measures are needed to prevent any loss to the marine aquaculture industry.

**Highlights:**

- This study reports detection of NNV in grouper larvae imported to Thailand
- The virus was assigned to the RGNNV genotype, which is the most extensively distributed genotype
- Histopathology revealed a pathognomonic lesion of NNV
- ISH using an NNV-specific probe revealed localization of NNV in the CNS and retina
- The virus was successfully propagated from diseased fish using the E-11 cell line

## 1. Introduction

Nervous necrosis virus (NNV) is the causative agent of viral nervous necrosis (VNN), also known as viral encephalopathy and retinopathy (VER). This virus is a member of the genus *Betanodavirus*, family *Nodavaridae* and has been responsible for disease outbreaks in both wild and farmed fishes (OIE 2019a; Bandín & Souto, 2020). Since the first incidence of VER or VNN reported over 30 years ago, more than 120 freshwater and marine fish species, including grouper (*Epinephelus* spp.), have been affected by NNV (OIE 2019a; Bandín & Souto, 2020; Xing et al., 2020).

NNV is a non-enveloped RNA virus with an icosahedral shape and a mean diameter of 25 to 30 nm. The NNV genome comprises two positively stranded RNAs (RNA1 and RNA2) (Breuil et al., 1991; Doan et al., 2017; Tan et al., 2015; Zorriehzahra, 2019). The RNA1 gene is 3.1 kb long and encodes the RNA-dependent RNA polymerase (RdRp) enzyme required for virus replication. RNA2 (1.4 kb) encodes the capsid protein of NNV. In virus-infected cells, RNA1 transcribes sub-genomic RNA3, which encodes proteins B1 and B2. The B1 protein inhibits necrotic cell death, thereby aiding virus replication. Simultaneously, the B2 protein protects the virus by inhibiting RNA splicing (Low et al., 2017).

Different genotypes of NNV have been identified based on the homology of RNA2, and their geographical distribution patterns were affected by water temperature (Bandín & Souto, 2020). Red-spotted grouper NNV (RGNNV), Tiger puffer NNV (TPNNV), Striped Jack NNV (SJNNV), and Bar-fin NNV were the first four genotypes identified (Nishizawa et al., 1995). New genotypes were discovered later, including the turbot betanodavirus strain (TNV) and Korean shellfish NNV (KSNNV) (Johansen et al., 2004; Kim et al., 2019). RGNNV is widely distributed among these genotypes, affecting the greatest number of fish (Bandín & Souto, 2020).

Pearl gentian grouper is cross-hybrid between giant grouper (*Epinephelus lanceolatus* ♂) and tiger grouper (*E. fuscoguttatus* ♀) (Jiang et al., 2015). This species inherits favourable characteristics from both parents, including disease resistance, rapid growth, and high-quality flesh; as a result, it is gaining popularity in China and Southeast Asian countries (Xing et al., 2020). In 2019, a disease outbreak killed 99% of imported pearl gentian grouper fry in a marine hatchery in Thailand. The infected larvae showed darkening, floating on the water surface, ‘sleepy behaviour’, and erratic swimming. In this study, we employed various diagnostic methods, including PCR, histopathology, *in situ* hybridization, genome sequencing and phylogenetic analysis, as well as cell culture to identify and characterize the imported causative virus associated with this unusual disease event.

## 2. Materials and methods

### 2.1 Fish sample collection

The samples used in this study were obtained from the imported pearl gentian grouper (*E. fuscoguttatus × E. lanceolatus*) fry from a marine hatchery in Thailand that experienced massive mortality of up to 99% in the quarantine tanks in August 2019 due to an unknown disease event. We received a collection of samples to identify the causative agent, including whole-body fry preserved in 10% neutral buffered formalin for histological examination and freshly dead fish for PCR diagnosis and cell culture.

### 2.2 Nucleic extraction and viral pathogen detection by PCR

Three pooled samples of diseased fish (3 individuals/pool) were subjected to RNA extraction using Trizol reagent (Invitrogen) according to manufacturer’s protocol and DNA extraction using lysis buffer containing 2% sodium dodecyl sulphate (SDS) and 1 μg/mL proteinase K prior to conventional phenol/chloroform extraction and ethanol precipitation as previously described (Meemetta et al., 2020). The quality and quantity of the obtained DNA and RNA were measured by spectrophotometry at OD 260 and 280 nm. Single PCR or RT-PCR assays were performed based on established protocols to diagnose NNV (OIE, 2019a), *Megalocytivirus* ISKNV (Gias et al., 2011), and *Ranavirus* (OIE, 2019b) from the obtained nucleic acid samples. 200 ng of either DNA or RNA template was used in a reaction. Quantification of NNV was later conducted using TaqMan probe-based qPCR according to the published method (Hick & Whittington, 2010). Table 1 shows a list of the PCR primers and probes used in this study. The pGEM T-easy (Promega) based positive control plasmid of NNV was constructed from a 605 bp RNA2 fragment amplified from NNV-infected orange-spotted grouper (laboratory sample). It was also used for building a standard curve for viral quantification in the RT-qPCR assay. Similarly, pGEM T-easy containing a 214 bp fragment of the *Megalocytivirus* ISKNV major capsid protein (MCP) gene (Dong et al., 2017) was used as a positive control for ISKNV detection. Note that this study did not have a positive control for *Ranavirus* detection.

**Table 1.**
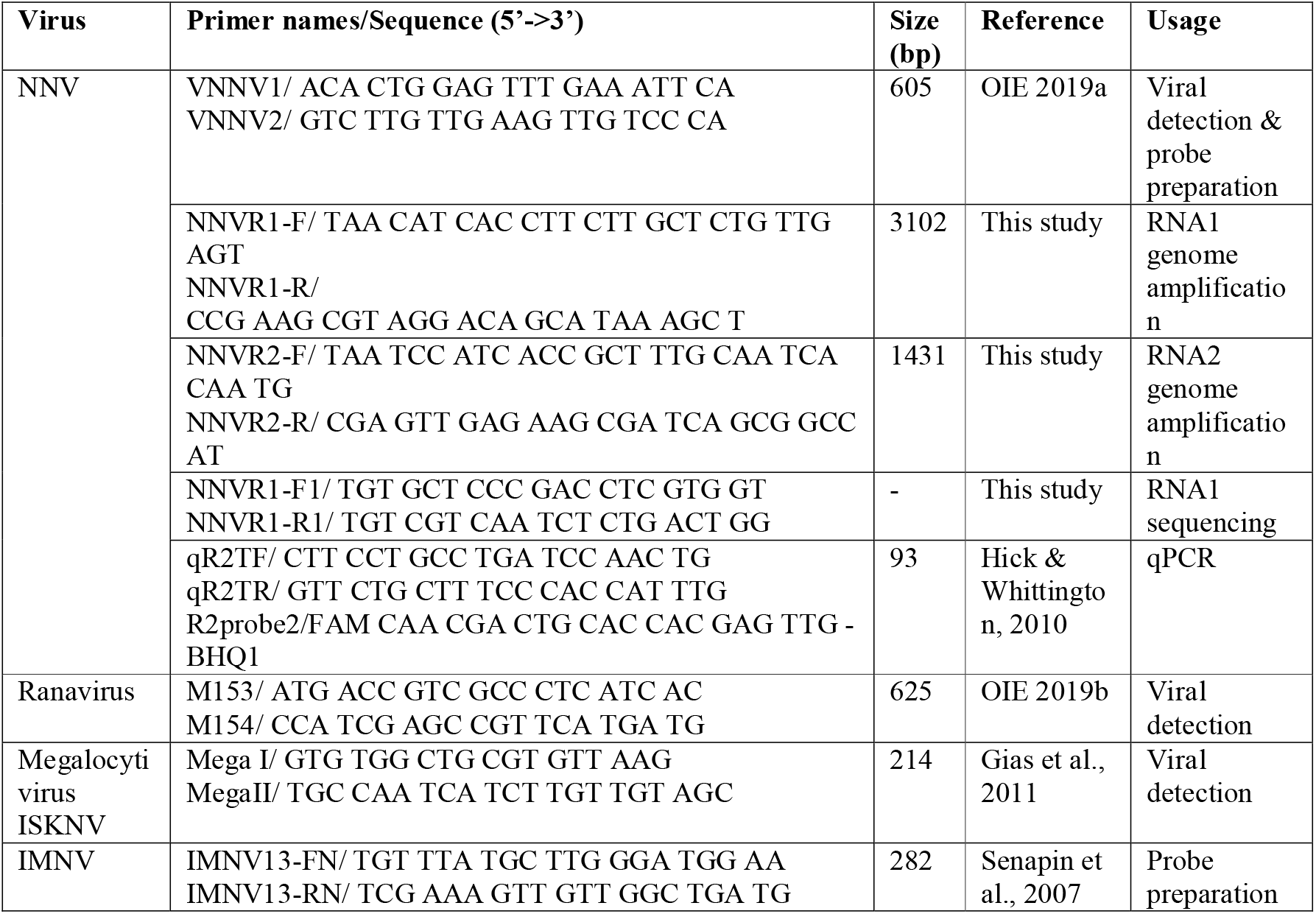
Primers used in this study.

### 2.3 Nervous necrosis virus genome amplification

Primer sets (Table 1) used for amplification of genomic RNAs of NNV were designed based on genome segments of red-spotted grouper nervous necrosis virus (RGNNV, accession numbers NC_008040 and NC_008041) and sevenband grouper nervous necrosis virus (SGNNV, accession numbers AB373028 and AB373029). RNA extracted from diseased grouper was used as a template for viral genomic segment amplification. RT-PCR reaction was carried out in a final volume of 25 μL containing 500 ng RNA template, 200 nM of each primer, 1 μL of SuperScript III RT/Platinum Taq Mix (Invitrogen), and 1 × supplied buffer. The amplification conditions were reverse transcription at 50 °C for 30 min and denaturation at 94 °C for 2 min followed by 30 cycles of 94 °C for 30 s, 60 °C for 30 s, 72 °C for 3 min (for RNA1) or 1.5 min (for RNA2), and final extension at 72 °C for 5 min. RT-PCR amplicons were electrophoresed on agarose gel, purified, and cloned into a pGEM-T easy vector (Promega). Recombinant clones were then sent to Macrogen (South Korea) for DNA sequencing using T7 and SP6 promoter primers. A recombinant clone of RNA1 was additionally read to fill the gap with NNVR1-F1 and NNVR1-R1 primers designed in this study (Table 1).

### 2.4 Phylogenetic tree construction

The amplified sequences of NNV from the diseased fish were subjected to homology searches using the NCBI blast tool. For phylogenetic analysis, publicly available genomes of NNV isolates whose bi-segmented genome belonged to the same genotype (not reassortment) were selected (Table 2) and aligned with the NNV sequences from this study. Following multiple sequence alignments and trimming, the phylogenetic trees were constructed based on the consensus sequences using Maximum Likelihood with the best model TN93+G suggested by MEGA10 software. Bootstraps was performed at 1,000 replicates.

**Table 2.**
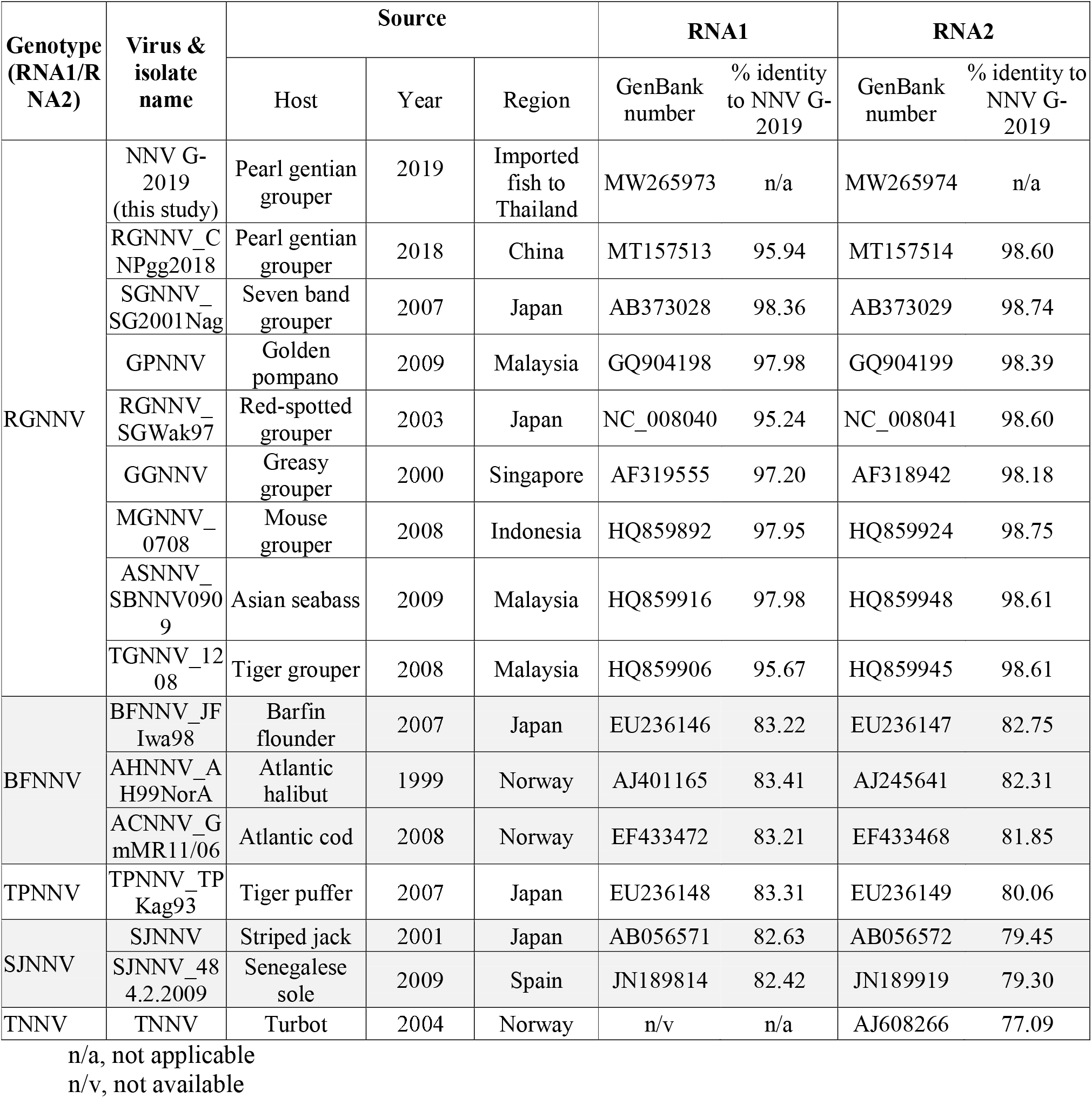
Piscine NNV sequences used for phylogenetic analysis

### 2.5 Histological examination and in situ hybridization (ISH)

The preserved specimens (n=5) were dehydrated and embedded in paraffin prior to consecutive sectioning for haematoxylin and eosin (H&E) staining and *in situ* hybridization (ISH) assay. An ISH assay using an NNV-specific probe was performed to determine the presence of the NNV viral genome in diseased fish tissues. Digoxygenin (DIG)-labelled probes were prepared using a commercial DIG-labelling mix (Roche, Germany). The recombinant plasmid containing a 605 bp insert derived from the NNV RNA2 fragment mentioned above was used as a template in labelling reaction, whereas a 282 bp fragment of infectious myonecrosis virus (IMNV) was used as an unrelated negative control probe (Senapin et al., 2007). The ISH procedure was carried out as previously described (Dinh-Hung et al., 2021) and a digital microscope was used to analyze and photograph the section.

### 2.6 NNV isolation using E-11 cell line

The tissue sample from NNV positive fish was homogenized in Leibovitz 15 (L-15) medium. The homogenized tissue was centrifuged at 10,000 g for 20 min at 4° C, and the suspension was filtered through a 0.22 μm membrane filter. The collected filtrate was used for infection of the E-11 cell line. The procedure for cell culture was performed as previously described by (Yamashita et al., 2005) with minor modifications. Briefly, E-11 cell lines were grown in L-15 medium supplemented with 5% fetal bovine serum (FBS). The monolayer of the E-11 cell line was inoculated with 500 μL of filtrate at 27 C for 1 h and then replaced by a new culture medium. The control E-11 cell line was inoculated with L-15 medium only. The cytopathic effect (CPE) was observed daily using an inverted microscope. For confirmation, three continuous passages were performed on the E-11 cell line for CPE observation, and culture supernatant was subjected to RNA extraction and RT-qPCR (Hick & Whittington, 2010).

## 3. Results

### 3.1 Clinically sick fish were tested positive for NNV

Three pooled samples of the diseased fish were tested positive for NNV by a single RT-PCR assay, indicating severe infection (Fig. 1A). On the contrary, these samples were tested negative for *Megalocytivirus*, ISKNV and *Ranavirus* as no amplification was detected (Fig. 1B, C). The result obtained from RT-qPCR showed that NNV positive fish samples had high virus loads ranging from 8.08 ×10^6^ to 2.64 ×10^7^ viral copies/200 ng RNA template (Fig. S1).

**Fig. 1.**
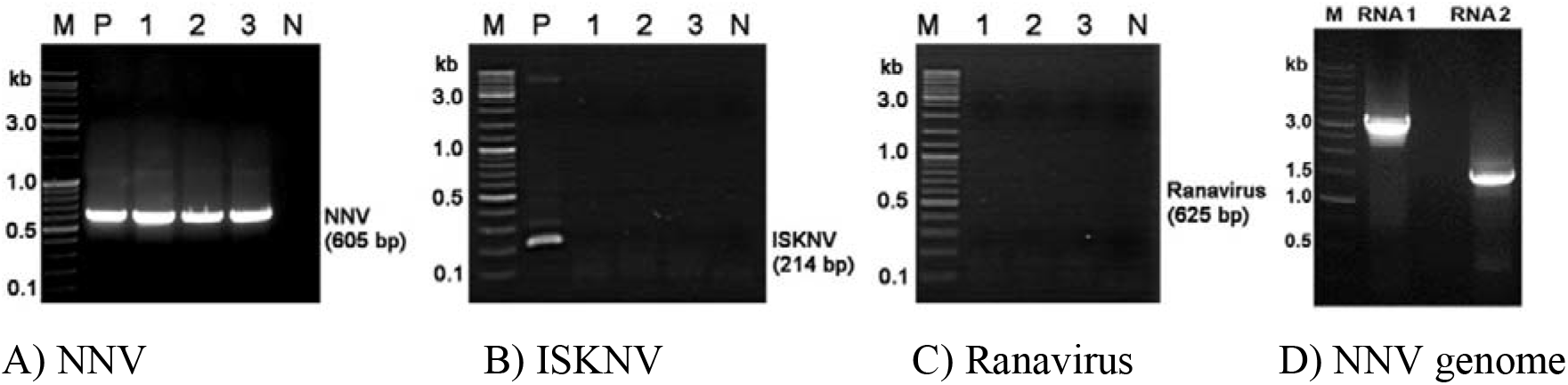
The photomicrographs of agarose gel electrophoresis after PCR amplification of NNV (A), Megalocytivirus ISKNV(B), and Ranavirus (C), NNV RNA1 (3,102 bp) and NNV RNA2 (1,431 bp) (D). M, marker; P, positive control; N, no template control; 1-3, tested pooled samples.

### 3.2 The genome sequence revealed RGNNV infection

The whole genome of NNV was obtained by amplifying RNA1 and RNA2 using their specific primers. Two expected bands of 3,102 bp (RNA1) and 1,431 bp (RNA2) were successfully amplified from RNA extracted from diseased fish and sequenced (Fig. 1C). The RNA1 and RNA2 nucleotide sequences were deposited to the NCBI GenBank database under accession numbers MW265973 and MW265974, respectively. The viral isolate in this study was named NNV G-2019. The RNA1 of NNV G-2019 predictably contained two ORFs encoding an RdRp protein (982 amino acids) and a protein B (75 amino acids). The RNA2 putatively encoded a capsid protein, which has 338 amino acids.

The blast results showed that nucleotide sequences of NNV G-2019 had significantly high percent identity to the represented viruses belonging to RGNNV genotype (95.24-98.36% for RNA1 and 98.18-98.74% for RNA2, Table 2) while lower percent identity to other genotypes (77.09-83.41%, Table 2). Moreover, the phylogenetic trees generated using representative genome sequences of five genotypes of NNV revealed that the genome sequences of NNV G-2019 were consistently clustered in the group of RGNNV genotype based on both RNA1 and RNA2 sequences (Fig. 2A-B). Notably, the genome identities of the viral isolates isolated from pearl gentian grouper in this study and a previous study (Xing et al., 2020) were 95.94% for RNA1 and 98.60% for RNA2 (Table 2) (Xing et al., 2020)

**Fig. 2.**
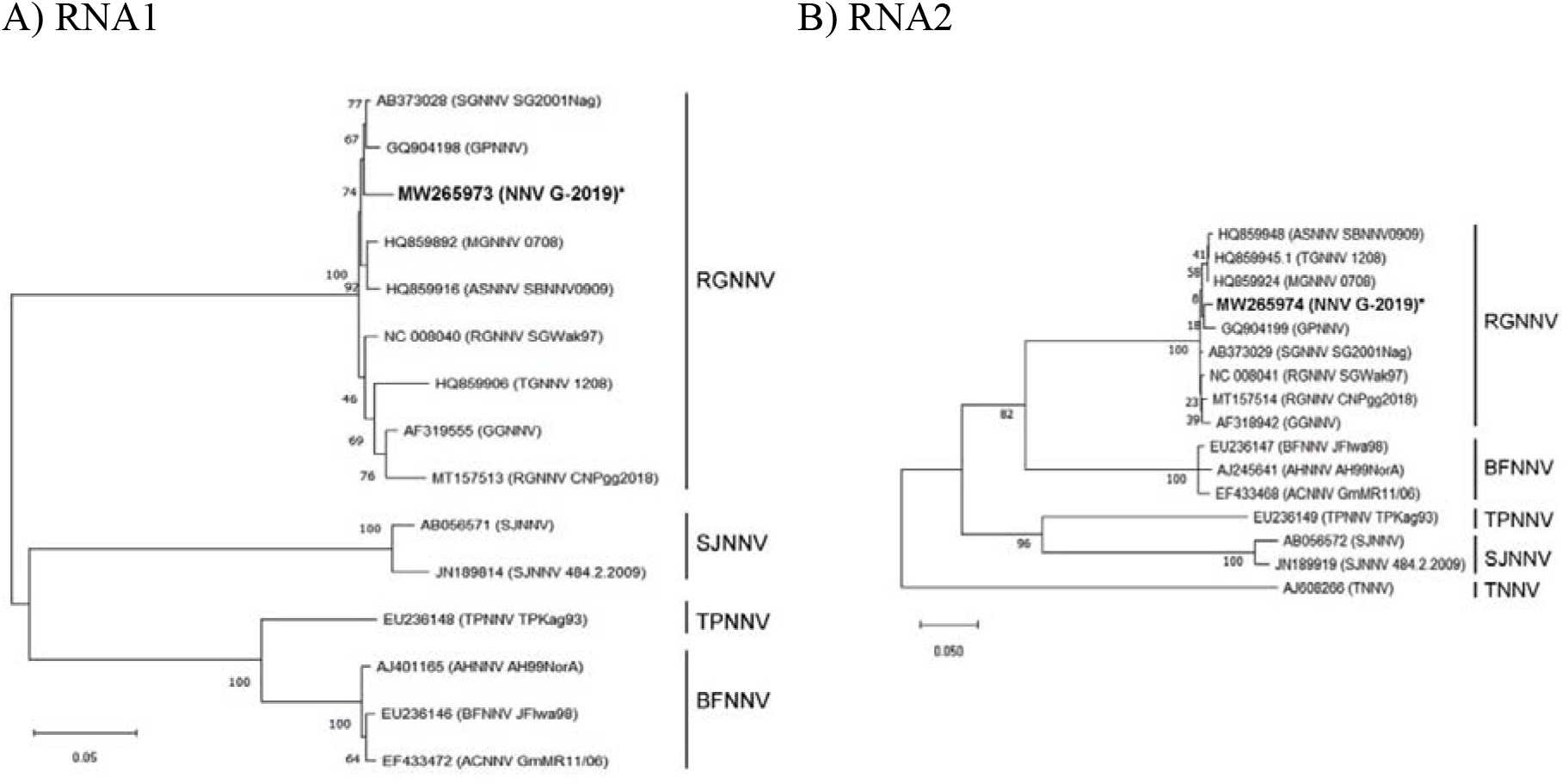
The maximum likelihood phylogenetic trees were constructed using NNV RNA1 (A) and RNA2 sequence (B) from this study and publicly available sequences obtained from the NCBI database. Consensus lengths for RNA1 and RNA2 were 2841 and 981 nucleotides, respectively. Bootstraps of 1,000 replicates were performed. * Denotes NNV G-2019 isolate obtained from this study.

Neuronal vacuolation, a pathognomonic lesion of NNV infection, was obviously observed in the outer and inner nuclear layers of the retina, the spinal cord, and occasionally observed in the brain of clinically sick fish (Fig. 3, left panel). ISH using an NNV-specific probe, detected dense positive signals in the retina and central nervous system (brain and spinal cord) of diseased fish (Fig. 3, middle panel). The ISH assay performed on the same samples using an unrelated probe (IMNV) revealed no hybridization signals (Fig. 3, right panel).

**Fig. 3.**
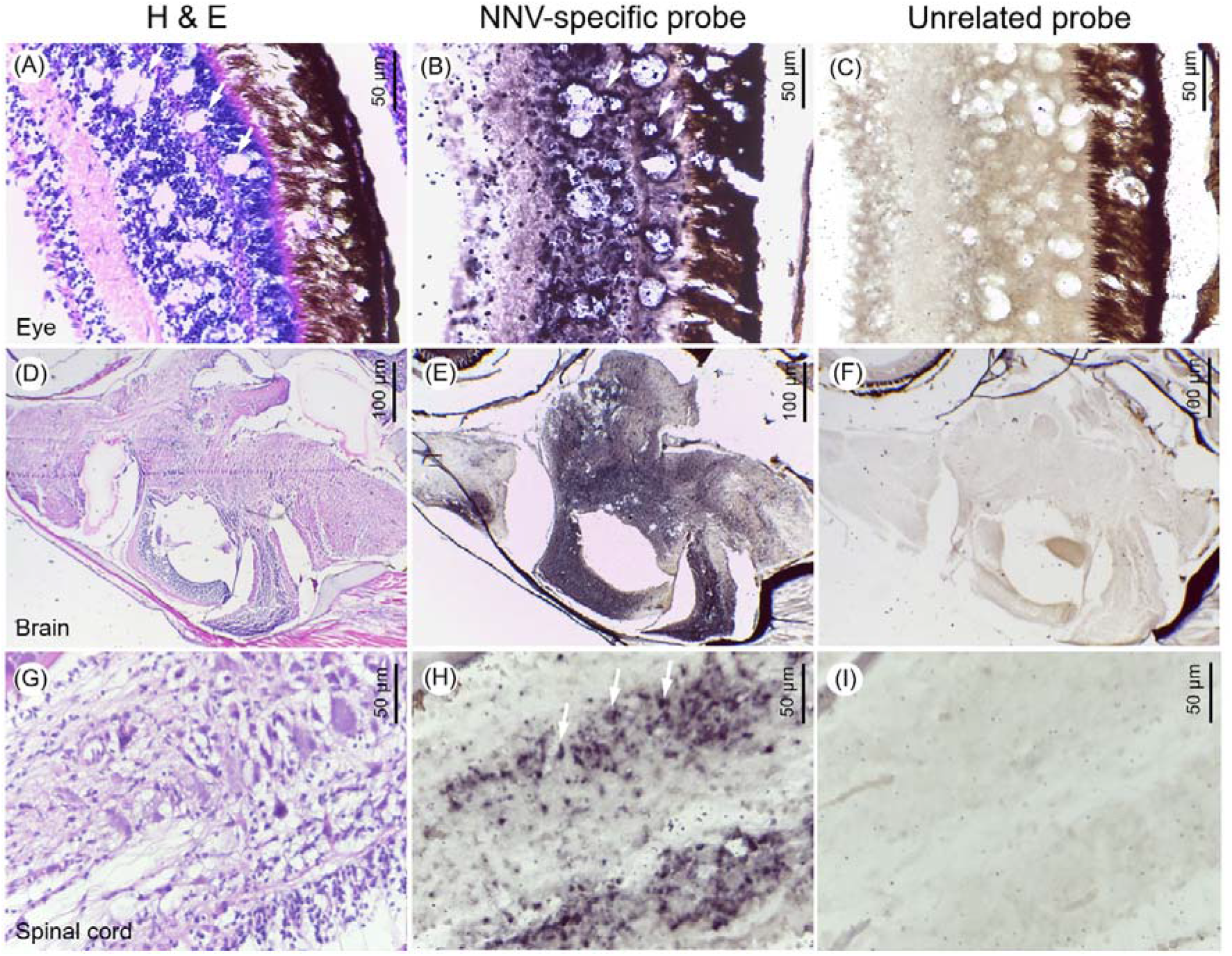
Consecutive sections of the eye (A-C), brain (D-F) and spinal cord (G-I) of grouper infected with NNV subjected to H&E-staining, ISH using NNV-specific probe and unrelated probe, respectively. Arrows in H&E-stained section indicate vacuoles. Arrows in ISH using NNV probe demonstrate positive signals. No signals were detected using an unrelated probe (C, F and I).

### 3.4 RGNNV infection revealed by cell culture method

Under the light microscope, cytopathic effect (CPE) was observed in the cells transfected with filtrate of diseased fish at 3 days post-inoculation (dpi). The main characteristic of CPE was intracellular vacuolation. At 5 dpi, clearer CPE was observed, including increased vacuoles, rounded cells and even detachment of cells from the flask’s bottom (Fig. 4). No CPE was observed from control cells. In the RT-qPCR quantification of NNV from cell culture, supernatant from the third passage revealed high copy numbers of the virus, ranging from 1.66 × 10^8^ to 1.98 × 10^9^ copies/mL (Fig. S1). No viral copies were detected from control cells.

**Fig. 4.**
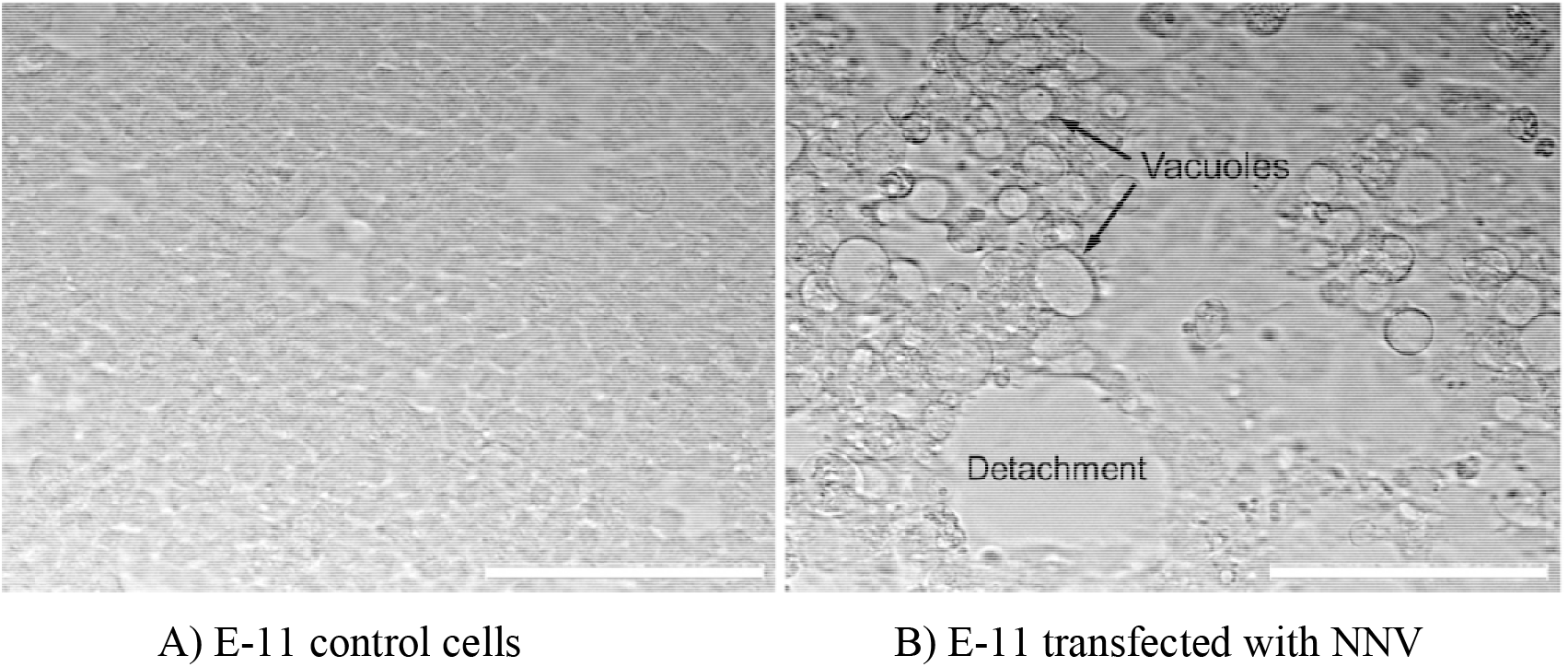
The CPE including multiple vacuoles, cell rounded and detachment observed from the E-1l cell line inoculated with filtrate of NNV-infected fish for 5 days at 25 C (B). No CPE was observed from the control cell line (A). Bar, 100 μm.

## 4. Discussion

The NNV is a highly contagious virus that has spread throughout the world. This virus has caused disease outbreaks in grouper hatcheries in Taiwan, Japan, China and other countries (Chi et al., 1997; Fukuda et al., 1996; Xing et al., 2020). Earlier in Thailand, there were some case reports of disease outbreaks caused by NNV in red-spotted grouper (*E. coioides*) and Nile tilapia (*Oreochromis niloticus*) larvae (Keawcharoen et al., 2015; Roongkamnertwongsa et al., 2005). The detection of NNV in pearl gentian grouper fry imported to Thailand highlights the critical role of improved biosecurity measures, particularly diagnostic testing and quarantine inspection, in mitigating the risk of disease spread from infected imported stock.

In this investigation, a combination of various disease diagnosis methods was applied for confirmation of NNV infection in pearl gentian grouper fry. The phylogenetic analysis based on RNA1 and RNA2 consistently indicated that the imported NNV to Thailand belonged to the RGNNV genotype and that no reassortment event occurred in the genome of this strain (called NNV G-2019). Previous reports of RGNNV in fish from Asian countries included the infections detected in Asian seabass from India (Parameswaran et al., 2008) and Malaysia (Ransangan & Manin, 2010), red-spotted grouper from Japan (Mori et al., 1991), grouper from Singapore (Hegde et al., 2002), and pearl gentian grouper from China (Xing et al., 2020). Although the sources of imported contamination were not specified, the viral genotype likely implied their possible origin from the same region.

As previously reported, an NNV outbreak in pearl gentian grouper juveniles in China resulted in 98% mortality and typical vacuolation of the retina and CNS (Xing et al., 2020). Neuronal vacuolation in the CNS, particularly the medulla oblongata, would result in abnormal swimming and neurological disorders (Xing et al., 2020). Our study further revealed the widespread presence of NNV throughout the brain, as indicated by dense ISH signals, supporting the described abnormal clinical symptoms of the NNV infected fish. NNV appears to have a significant impact on larvae and juveniles (Zorriehzahra, 2019; Xing et al., 2020), whereas adults remain an asymptomatic carrier of the disease (Gomez et al., 2004). Vertical transmission of NNV was reported in various fish species such as European seabass, seven-band grouper, Atlantic halibut (Breuil & Romestand, 1999; Grotmoll et al., 1999; Watanabel et al., 2000; Bandín & Souto, 2020) and was suspected in the outbreak case of pearl gentian grouper in China (Xing et al., 2020). Although the infected fry in this study could have been another case of vertical transmission, horizontal transmission of NNV through contaminated water, equipment, and feed cannot be ruled out. While NNV could be isolated in various cell lines, E-11 was found to be the most permissive cell line for *in vitro* propagation of NNV, including RGNNV, SJNNV, and reassortant RGNNV/SJNNV genotypes (Valero et al., 2021). In this study, the imported NNV isolate, NNV G-2019, was successfully propagated in the E-11 cell line and bio-banked. This is particularly useful for future research on the virus’s epidemiology and evolution, as well as for developing control measures.

In conclusion, this study reported a case of disease outbreak caused by NNV RGNNV genotype in a population of pearl gentian grouper fry imported to Thailand in 2019. The virus was successfully isolated and bio-banked for further investigation. Given the NNV’s diverse host specificity, which includes tilapia and Asian seabass, on which Thailand’s aquaculture industry thrives, adequate precaution and biosecurity practices are essential to avoid any losses.

## Acknowledgements

This study was supported by Mahidol University (Fundamental Fund: Basic Research Fund: fiscal year 2022, Grant number BRF1-054/2565). H.T.D acknowledges the Research Initiation Grant from the Asian Institute of Technology (AIT). S.V.J. was supported by HM Queen’s Scholarship from AIT. The authors thank Pimchanok Kansri and Sukhontip Sriisan for their skilled technical assistance.

## Credit authorship contribution statement

S.S. and H.T.D., conceptualization, methodology and supervision; S.V.J., S.T., P.K., K.P., investigation, data analysis; S.S., H.T.D and S.V.J., Writing original draft; M.S., B.N., and K.R.S., resources, review-editing. All authors have read and agreed to the current version of the manuscript.

## Supplementary data

**Fig. S1.**
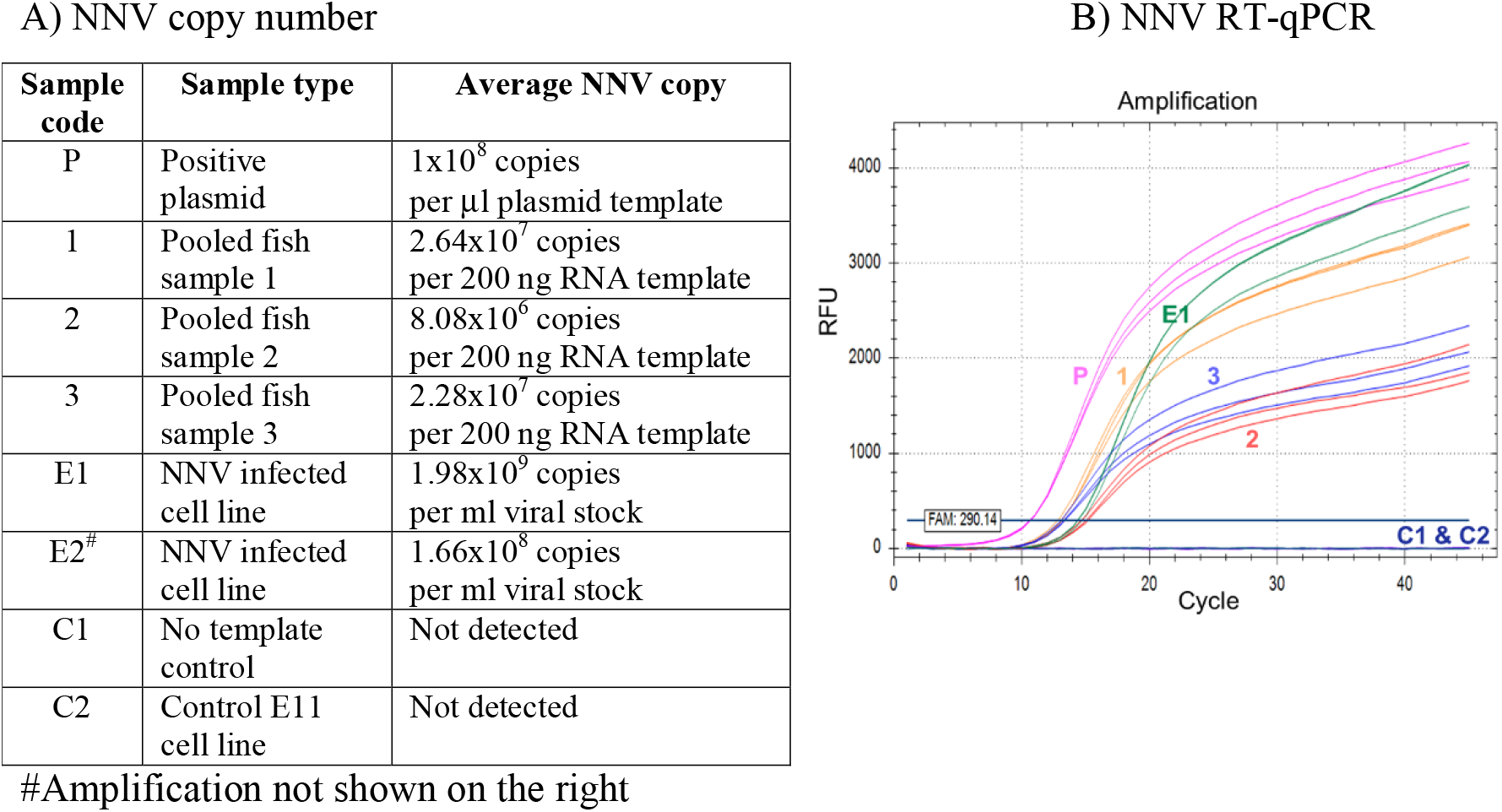
The viral quantification was done through the RT-qPCR, showing high viral loads from the sample of diseased fish (pooled sample numbers 1 to 3) and cell culture (E1-E2). P, Positive plasmid; C1, no template control; C2, control cell culture sample. Each sample was quantified in three replicate wells.

